# Newer Surveillance Data Extends our Understanding of the Niche of *Rickettsia montanensis* (Rickettsiales: Rickettsiaceae) Infection of the American Dog Tick (Acari: Ixodidae) in the United States

**DOI:** 10.1101/2023.01.11.523628

**Authors:** Catherine A. Lippi, Holly D. Gaff, Robyn M. Nadolny, Sadie J. Ryan

## Abstract

**Background:** Understanding the geographic distribution of *Rickettsia montanensis* infections in *Dermacentor variabilis* is important for tick-borne disease management in the United States, as both a tick-borne agent of interest and a potential confounder in surveillance of other rickettsial diseases. Two previous studies modeled niche suitability for *D. variabilis* with and without *R. montanensis*, from 2002-2012, indicating that the *D. variabilis* niche overestimates the infected niche. This study updates these, adding data since 2012.

**Methods:** Newer surveillance and testing data were used to update Species Distribution Models (SDMs) of *D. variabilis*, and *R. montanensis* infected *D. variabilis*, in the United States. Using random forest (RF) models, found to perform best in previous work, we updated the SDMs and compared them with prior results. Warren’s I niche overlap metric was used to compare between predicted suitability for all ticks and ‘pathogen positive niche’ models across datasets.

**Results:** Warren’s I indicated <2% change in predicted niche, and there was no change in order of importance of environmental predictors, for *D. variabilis* or *R. montanensis* positive niche. The updated *D. variabilis* niche model overpredicted suitability compared to the updated *R. montanensis* positive niche in key peripheral parts of the range, but slightly underpredicted through the northern and midwestern parts of the range. This reinforces previous findings of a more constrained pathogen-positive niche than predicted by *D. variabilis* records alone.

**Conclusions:** The consistency of predicted niche suitability for *D. variabilis* in the United States, with the addition of nearly a decade of new data, corroborates this is a species with generalist habitat requirements. Yet a slight shift in updated niche distribution, even of low suitability, included more southern areas, pointing to a need for continued and extended monitoring and surveillance. This further underscores the importance of revisiting vector and vector-borne disease distribution maps.

## Introduction

Species distribution models (SDMs) are increasingly utilized to estimate the geographic distribution of infectious diseases, particularly those caused by agents transmitted by arthropod vectors. The basic methodology for constructing SDMs (or ecological niche models) consists of combining species occurrence data with continuous layers of environmental predictor variables, which are fed into a modeling algorithm (Elith and Franklin, 2013; Franklin, 2010; Peterson and Soberón, 2012). The resulting model is projected onto a defined study area, yielding spatially continuous habitat suitability estimates for areas of the landscape that were not originally sampled. Species distribution modeling is an intuitive approach to delineating vector-borne disease ranges that is logistically feasible, particularly when surveillance programs or capacity for pathogen testing are limited. When faced with multiple unknowns (e.g., unknown transmission cycles, emerging novel pathogens, etc.), the distribution of vectors on the landscape are sometimes used in a public health context to approximate risk of exposure to pathogens (Lippi et al., 2021b, 2021c). Yet, it is important to differentiate between the distribution of the vectors and that of the pathogens they transmit. Vector presence is not in itself sufficient for pathogen transmission to occur. Precise delineation of geographic risk facilitates the development of targeted health policies, educational campaigns, and interventions with the potential to avert the misallocation of limited resources.

The need for geographically conservative assessments of transmission risk is perhaps most evident with cosmopolitan vectors, whose broad geographic ranges may far exceed the limits of known transmission to humans. The American dog tick (*Dermacentor variabilis*) is a medically important arthropod vector of several zoonotic pathogens, including the causative agents of Rocky Mountain spotted fever (RMSF) (*Rickettsia rickettsii*) (Brumpt; Rickettsiales: Rickettsiaceae) and tularemia (*Francisella tularensis*) (Dorofe’ev; McCoy and Chapin; Thiotrichales: Francisellaceae). Both of these diseases can be fatal without medical intervention, perhaps justifying medical advisories that equate risk of tick exposure with transmission risk, particularly when surveillance data are scarce, or in cases where ticks themselves act as reservoir hosts (CDC, 2022). In addition to RMSF, *D. variabilis* also transmits other spotted fever group (SFG) rickettsial agents, as well as *R. montanensis* (Rickettsiales: Rickettsiaceae), a rickettsial group agent that is suspected of causing nonfebrile rashes in humans, and has caused clinical symptoms in an animal model (McQuiston et al. 2012; Snellgrove et al. 2021). Although not included in the case definition for SFG pathogens, it is likely that *R. montanensis* infections may account for some of the recent increases in SFG reporting, as immunological cross-reactivity between rickettsial pathogens is frequently observed with commonly used serologic tests (Abdad et al. 2018). Of note, *D. variabilis* has recently been proposed to be split into two species, with a western portion of the population as a distinct species, *D. similis* (Lado et al., 2021); however, we do not differentiate in this study.

Determining the geographic risk of *D. variabilis* infection with *R. montanensis* has profound implications for the management of tick-borne diseases in the United States, as both a tick-borne agent of interest and a potential confounder in the surveillance of other Rickettsial diseases. A model of the distribution of *D. variabilis* and *R. montanensis* positive samples was published by St John et al. in 2016, using MaxEnt modeling to describe and predict environmental suitability in the United States, based on data obtained through the Department of Defense (DoD) Human Tick Test Kit Program, now called the Military Tick Identification/Infection Confirmation Kit Program (MilTICK). These data were available at the time through the VectorMap online data platform (http://vectormap.si.edu/dataportal/) (St John et al., 2016). The MilTICK data were human-biting ticks submitted from U.S. military installations as part of a tick-testing program; test results were reported back to the bitten individuals, and the data were also used as passive vector surveillance. In 2021, Lippi et al. re-examined the distribution of *D. variabilis* and the *R. montanensis* infected niche in the USA, both to understand whether predicted risk of suitability for tick encounters or infected tick encounters were distinct, and to explore and compare multiple modeling approaches for assessing the distribution of this tick vector (Lippi et al., 2021a). The 2021 study was able to leverage the original dataset used in the 2016 study, and used a refined set of environmental predictors to compare a suite of Species Distribution Model (SDM) approaches. Lippi et al. found support for an “infected niche” within the broader distribution of *D. variabilis* which was largely consistent across models, though the Random Forests (RF) approach (Breiman, 2001) provided the best performing models, given the available data (Lippi et al., 2021a). Though somewhat limited in terms of the full geographic distribution of *D. variabilis* ticks (i.e., few locations were reported from the tick’s southern extent), the dataset used in these studies provided a rare opportunity to directly assess the distribution of pathogens within vectors, as every individual tick collected had been tested for *R. montanensis* as part of an extensive passive surveillance network. Both of these studies demonstrated that *D. variabilis* ticks infected with *R. montanensis* had estimated geographic distributions that were considerably restricted compared to that of *D. variabilis* alone, thus supporting an “infected niche” that exists as a subset of the vector’s full range.

In the current study, we revise the *D. variabilis* distribution maps using occurrence data updated with novel surveillance points collected since 2012, and further refine the environmental variables according to current best practices using the RF approach (Escobar et al., 2014; Valavi et al., 2021). We explore whether the additional data impact the estimated suitability distribution, the relative importance of environmental input variables, and mapped prediction outputs.

## Methods

*Tick Surveillance Data* – Two previous studies on *D. variabilis* in the United States were conducted using occurrence locations recorded in the continental United States from 2002 to 2012, where ticks were tested for *R. montanensis* as part of MilTICK, and are described in St John et al. (2016) and Lippi et al. (2021) (Lippi et al., 2021a; St John et al., 2016).

Georeferenced data were openly available through VectorMap (http://vectormap.si.edu/dataportal/), a project of the Walter Reed Bioinformatics Unit (WRBU), housed at the Smithsonian Institution Washington DC (St John et al., 2016). All ticks submitted through MilTICK are tested for rickettsial pathogens via PCR as previously described (Milholland et. al., 2021, Stromdahl et al., 2011), providing information on infection status (i.e., true presence or absence) for the entire dataset. Exposure locations were determined by asking MilTICK participants to self-report where the tick bite was most likely acquired, accounting for travel history. If no separate information on tick-bite location was submitted, ticks were assumed to be acquired on or near the military installation from which the tick was submitted.

New records of *D. variabilis* reported and tested for *R. montanensis* through MilTICK since 2012 through 2021 were made available for this study. These data were de-identified, and though general locality data were provided (e.g., military installation where reported, or towns and cities where ticks were collected), positional coordinates were not provided. New surveillance data were manually georeferenced for this study, following the general protocol reported in the metadata of the original dataset (i.e., 2002-2012 records) georeferenced for TickMap by the WRBU. Geographic coordinates (i.e., latitude and longitude) were assigned to records, taking the centroid of named locations found in Google Maps. Spatial uncertainty for points was established based on the spatial extent of reported locations (e.g., municipal boundaries, reported area of military installations, etc.). We excluded records where the spatial uncertainty exceeded 10km, ensuring that the spatial resolution of the St. John et al. (2016) and Lippi et al. (2021) studies was matched for all analyses.

We removed duplicate records and records without pathogen testing results (n=14). Data thinning on the remaining species occurrence points was performed via the ‘spThin’ package in R (ver. 4.1.2) (R Core Team 2019), which uses a spatial thinning algorithm to randomly remove excess occurrence locations within a specified distance threshold (Aiello-Lammens et al., 2015). This was performed for both the original data in the Lippi et al. 2021 study and the updated dataset to reduce susceptibility to geographic sampling bias, for example, when overrepresented locations erroneously drive species environmental associations due to repeated observations at discrete locations. Due to the passive nature of the tick surveillance program, it was deemed necessary to thin occurrences and minimize the potential effect of sampling bias, where locations near medical facilities and military installations may be inherently overrepresented. This process resulted in one unique, randomly selected location per 10km, and was performed on the full dataset of tick records, and on the subset of ticks that tested positive for *R. montanensis*.

The original dataset used to build the distribution models reported in Lippi et al. 2021 was then compared to an updated dataset, reflecting new surveillance data. Because new surveillance data consisted of fewer records compared to the original study, the updated dataset was comprised of both original surveillance data and new surveillance records. Following the framework of Lippi et al. 2021, we estimated separate geographic distributions of *D. variabilis*, and the subset of records that tested positive for *R. montanensis* infections, for both the original and updated tick surveillance records. Environmental data layers used in modeling consisted of interpolated bioclimatic (bioclim) layers from WorldClim (ver. 2), and gridded soil variables (0cm standard depth) taken from International Soil Reference Information Centre (ISRIC) SoilGrids (Fick and Hijmans, 2017; Hengl et al., 2017). Gridded environmental data inputs were used at 10km resolution to match the scale of tick occurrence data. Bioclim layers with known errors (i.e., Bio8, Bio9, Bio18, and Bio19) were removed a priori, and Variance Inflation Factor (VIF) was used to control for collinearity in the remaining variables (th=10) (Escobar et al., 2014). The final set of variables used to build models included annual mean temperature (Bio1), mean diurnal range (Bio2), temperature seasonality (Bio4), precipitation of wettest month (Bio13), precipitation of driest month (Bio14), precipitation seasonality (Bio15), soil organic carbon density (OCDENS), available soil water capacity until wilting point (WWP), and soil pH (PHIHOX).

Random forests (RF) modeling, implemented in R with the package ‘sdm’, was used to estimate tick distributions, following recommendations for settings and parameters described in Valavi et al 2021 (Valavi et al., 2021). We ran 500 RF model replicates for each dataset of occurrence points (i.e., original and updated records for all *D. variabilis*, and original and updated records for only *D. variabilis* infected with *R. montanensis*), averaging projected model output to produce four estimated distributions. Average model accuracy metrics for each experiment were calculated to assess the predictive accuracy of SDMs against a random holdout of 25% data from each dataset, respectively. Four measures were calculated to assess model accuracy, the receiver operator characteristic (ROC) curve with area under the curve (AUC), true skill statistic (TSS), model deviance, and mean omission (i.e., false negatives). We quantified the niche overlap between averaged models with the Warren’s I index, calculated in R with the package ‘spatialEco’ (Warren et al., 2008). The I statistic is an indicator of the similarity between two distributions, with values ranging from 0 (i.e., no overlap in the niche) to 1 (i.e., the niche is identical). A difference map to assess agreement in suitability predictions between the updated full dataset and infected dataset models was generated in R using the packages Raster and RasterVis by taking the difference of model output rasters and plotting them.

## Results

Updated input surveillance data increased our sample sizes for the full dataset (original n=432, updated n=525), and for the ticks positively identified for *R. montanensis* infection (original n=44, updated n=63). We found that updating the input data increased the spatial extent of predicted suitability for both the full dataset of all ticks (Figure 1 A (original) and B (updated)) and for the infected dataset (Figure 1 C (original) and D (updated)). Although we made no distinction for potential records of the newly described species *D. similis*, a few occurrence points were from the Western United States (original n=10, updated n=21). Model accuracy metrics for averaged RF models across the four datasets are presented in Table 1. Accuracy metrics across models indicated generally good performance, with AUC values exceeding 0.90, and TSS values greater than 0.64. Though comparable in output, averaged models made with updated data performed lower than models made with original datasets, indicated by lower AUC and TSS values, and higher deviance and omission. A Warren’s I index comparison of the original and updated dataset suitability predictions for the full and infected niche, showed they differed by less than 2% each (full dataset: full dataset =0.981, positive dataset: positive dataset =0.986).

**Table 1.** Average model accuracy metrics for Random Forest models, using different datasets of tick occurrences.

| Dataset | Subset | AUC | Deviance | TSS | Omission |
| --- | --- | --- | --- | --- | --- |
| Original* | All Ticks | 0.953 | 0.570 | 0.769 | 0.116 |
| Original* | Positive Ticks | 0.930 | 0.690 | 0.710 | 0.145 |
| Updated | All Ticks | 0.918 | 0.742 | 0.692 | 0.154 |
| Updated | Positive Ticks | 0.905 | 0.812 | 0.643 | 0.179 |

\*data used in Lippi et al. 2021

**Figure 1:**
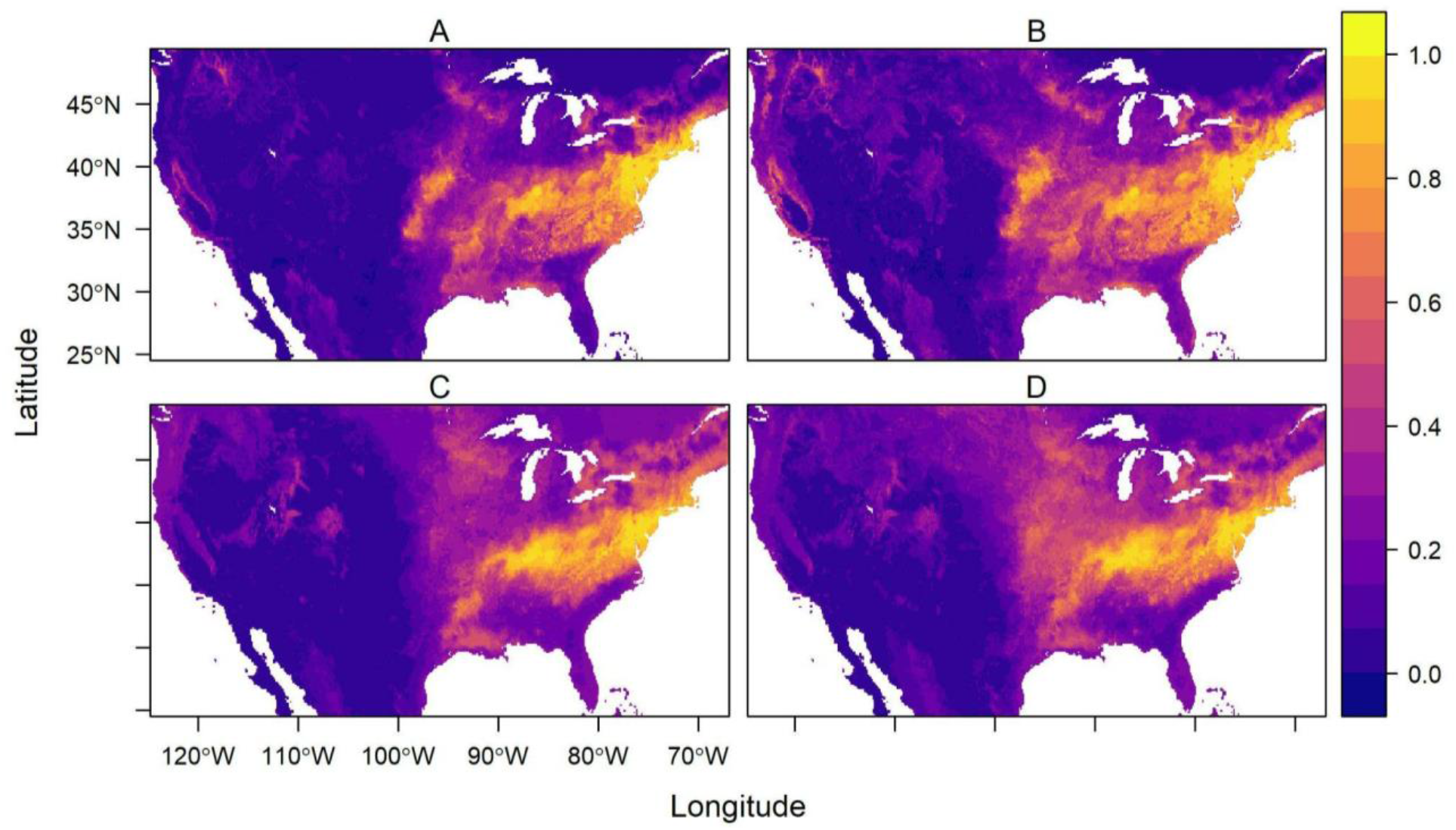
Predicted habitat suitability from average output of 500 random forest models for the original (A, C) and updated (B, D) datasets for all *D. variabilis* data (A, B), and *D. variabilis* infected with *R. montanensis* (C, D)

The updated *R. montanensis* positive ticks, as in the original analyses, are predicted to have a niche which is a subset of the full predicted niche (Figure 1D). The Warren’s I comparisons of the ‘infected niche’ and the full datasets for original (full:infected =0.950), and updated datasets (full:infected = 0.968) suggest that these are not dissimilar predicted niche distributions where they overlap, yet they are not capturing identical distributions.

The importance of variables underlying model predictions varied across datasets, although precipitation seasonality (Bio15) was the top contributing environmental predictor in all models (Fig. 2). Mean diurnal range (Bio2) and precipitation of driest month (Bio14) were also relatively important variables in models of both the original and updated full tick datasets, though these variables did not contribute highly to the models of infected tick distributions.

**Figure 2:**
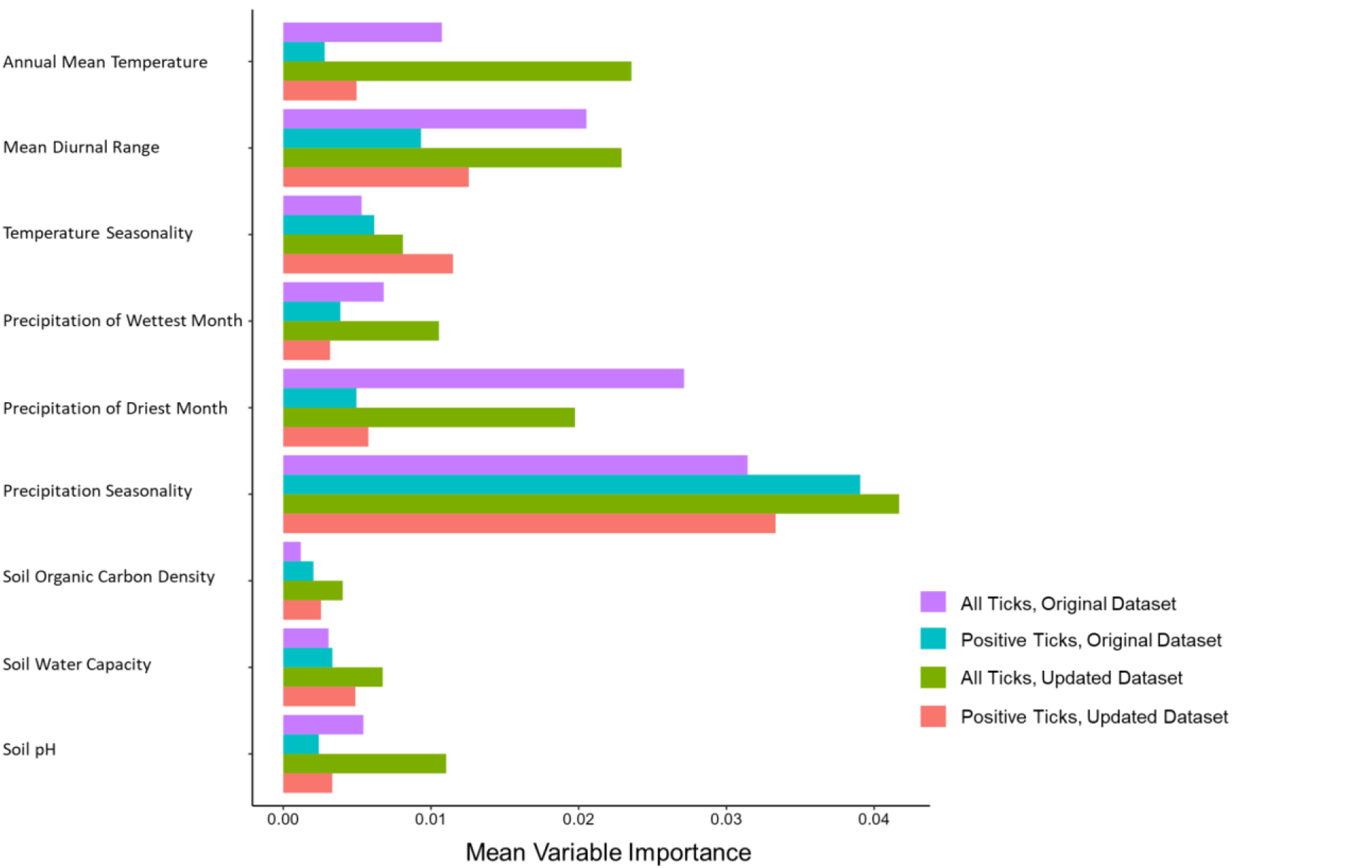
Relative variable importance from average output of 500 random forest models for the original and updated datasets for all *D. variabilis* data, and *D. variabilis* infected with *R. montanensis*.

To visualize the difference in predicted suitability for all ticks and that predicted for the pathogen-positive ticks, we visualized the difference in mapped suitability estimates from updated models (Fig. 3). The resulting map highlights the overprediction (redder colors) or underprediction (darker blue colors) of a model trained on all surveilled ticks, compared to one trained on *R. montanensis* positive ticks. Infected ticks are overpredicted by the model of all ticks along the southeastern and western peripheries of the infected tick distribution, and underpredicted to a lesser degree, along the northern border and through parts of the mid-Atlantic to midwestern states (Figure 3).

**Figure 3:**
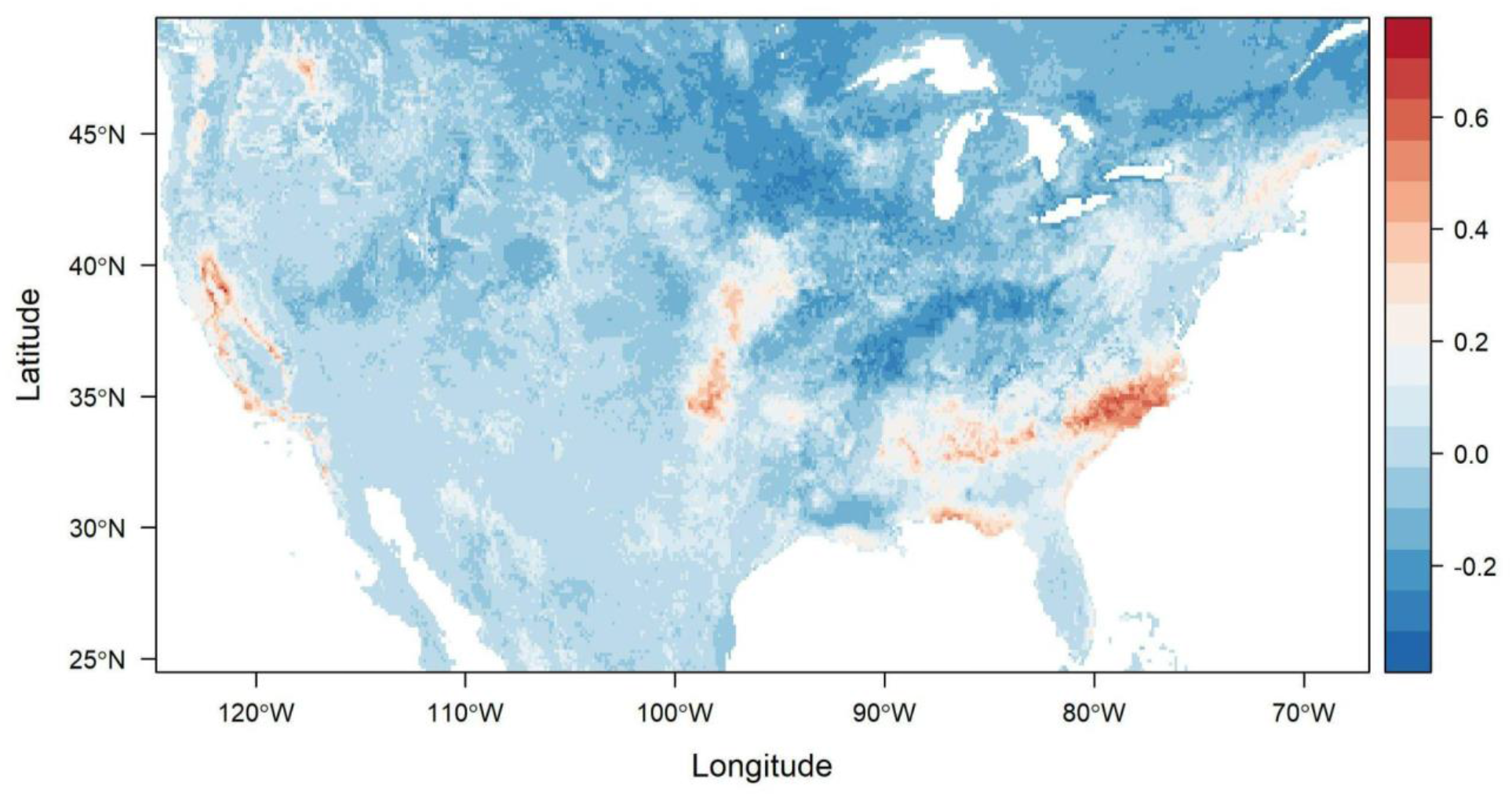
Assessing differences in predicted suitability for an average of 500 Random Forest models for *D. variabilis* and those infected with *R. montanesis* -redder colors depict overprediction by a tick-only model, and darker blue colors, underprediction.

## Discussion

A number of factors exist that influence SDM output, including sampling bias, choice of environmental predictors, modeling algorithm, and other user-specified inputs (Araújo et al., 2019; Valavi et al., 2021). In this study, we updated previously published RF models of *D. variabilis* and *D. variabilis* infected with *R. montanensis*. This update was made possible by the addition of surveillance and testing data to the original dataset used. We thus explored what impact the additional data had on predictions found previously, via modeling both datasets and comparing predicted suitability with a niche overlap metric, Warren’s I, and presenting the mapped output of modeled predictions using the original and updated datasets. We additionally presented a visualization of agreement, highlighting areas of over and underprediction of the infected niche by the overall niche prediction.

Models made with both datasets were generally high-performing, and overlap indices showed that suitability predictions varied only slightly with the inclusion of novel surveillance data. The estimated range of *D. variabilis* primarily extends throughout the eastern United States, with the highest predicted probabilities spanning areas in the Midwest, Mid-Atlantic, and Northeast regions. The southern boundary of *D. variabilis* occurrence was not well captured in Lippi et al. 2021, owing to limited data points from this region in the original MilTICK dataset. Although records of ticks from southern locations (e.g. Texas and peninsular Florida) exist in online repositories, these records were not included in efforts to directly compare distributions of ticks of known infection status. Notably, the predicted geographic distribution for *D. variabilis* extends further South in the updated model, indicated by higher probabilities of suitability in Texas and Florida.

The predicted suitability distribution of *D. variabilis* infected with *R. montanensis*, or infected niche, is geographically constrained, compared to the full predicted suitability distribution of *D. variabilis*, regardless of data inputs. Areas of range disagreement, highlighted by the difference map, are most prominent along the southern and western peripheries of the full *D. variabilis* range in the eastern US, as well as on the west coast. A potential explanation for this kind of pattern is that in the more established parts of the range -i.e. the more central parts of predicted range -there may be higher *R. montanensis* exposure risk. For different tick-borne pathogens, and even for different species of ticks, evidence of patterns of expansion by both the vector and the pathogen, together or temporally lagged have varied (Burrows et al., 2021; Dahlgren et al., 2016; Fornadel et al., 2011). This highlights the limitations inherent in using vector distribution maps as proxies for transmission risk maps directly; incorporating pathogen testing results into this type of distribution modeling can help constrain the area most likely to be important for disease transmission exposure risk. This is particularly germane for a generalist vector such as *D. variabilis*, where the presence of the pathogen in question may be patchily distributed. Disagreement along the West coast may also be influenced by the inclusion of *D. variabilis* records from California, Oregon, and Washington. The western population of *D*.*variabilis* has recently been proposed as a new species (*Dermacentor similis*), and thus may have fundamentally different habitat suitability requirements (Lado et al., 2021).

*Dermacentor* ticks are receiving increasing attention as significant vectors of zoonotic pathogens, and there have been recent calls for closer monitoring of understudied species (Lippi et al., 2021c; Martin et al., 2022). Species distribution modeling offers a framework for rapidly estimating potential distributions of vectors when ample occurrence data are available. Yet, there are considerable ramifications that may arise if models are put into public health practice without thorough assessment (Erdemir et al., 2020). It is therefore necessary to periodically review estimates of risk as new data or methods become available. However, in this study we found that an additional nine years of passive surveillance data resulted in negligible differences in distribution estimates. This points to the benefit of augmenting existing surveillance to target undersampled areas, and highlights the need to expand pathogen testing capabilities to other existing networks. Widespread, county-level surveillance for *D. variabilis* in the United States is currently limited (Lehane et al., 2019). Pathogens with low detection rates may particularly benefit from targeted, active surveillance strategies to delineate risk. In this study, updated passive surveillance data yielded only 19 novel spatially unique records of infected ticks after thinning. To contrast, a recent study that targeted a discrete area in Northern Wisconsin, an area of low predicted suitability in our models, successfully detected *R. montanensis* in *D. variabilis* (Vincent and Hulstrand, 2022). Focused testing efforts, particularly in locations bordering areas of range disagreement, may help resolve the limits of exposure risk and facilitate targeted monitoring efforts.

In conclusion, infected ticks are predicted to have a distribution that is a subset of the full vector range, a finding which is consistent across original and updated data inputs. For a generalist vector such as *D. variabilis*, ascertaining the key areas of pathogen exposure risk within such a large range of predicted suitability, is an important potential tool for future surveillance and monitoring. Revisiting the estimation of tick distributions is a necessary endeavor, particularly as we gain more information on tick-borne transmission cycles through surveillance and laboratory studies. There are few occurrence records that establish *D. variabilis* at the county level throughout our predicted suitability range in the contiguous United States, pointing to a general need for increased surveillance activities (Lehane et al., 2019). Yet, placing emphasis solely on new data collection for the refinement of spatial risk assessments may not yield dramatic gains in information. This is perhaps most evident in the passive surveillance of pathogens with low detection rates. Additionally, we suggest that there is a great need to validate the data in areas identified as high risk through active surveillance, particularly where passive surveillance is lacking. Moving forward, efforts to further refine geographic risk estimates of tick-borne pathogens will benefit from targeted surveillance to resolve distributional boundaries.

## Author Contributions

Catherine A. Lippi: Conceptualization, data duration, design of methodology, formal analysis, visualization, writing – original draft, writing – reviewing and editing. Holly D. Gaff: Conceptualization, writing – original draft, writing – reviewing and editing. Robyn M. Nadolny: Data curation, writing – original draft, writing – reviewing and editing. Sadie J. Ryan: Conceptualization, data curation, design of methodology, formal analysis, visualization, writing – original draft, writing – reviewing and editing

